# Infectious transgenesis in arthropods

**DOI:** 10.64898/2026.09.14.751565

**Authors:** Najmeh Nejat, Travis van Warmerdam, Jason L. Rasgon

**Affiliations:** Department of Entomology, The Pennsylvania State University, University Park, PA, USA; The Center for Infectious Disease Dynamics, The Pennsylvania State University, University Park, PA, USA; The Huck Institutes of the Life Sciences, The Pennsylvania State University, University Park, PA, USA; Department of Biochemistry and Molecular Biology, The Pennsylvania State University, University Park, PA, USA; One Health Microbiome Center, The Pennsylvania State University, University Park, PA, USA

**Keywords:** Baculovirus, AcMNPV, CRISPR, GEValT, ReMOT, mosquito, *Aedes aegypti*, *Anopheles*, *Culex*

## Abstract

In the last decade, there has been an explosion in the development of in vivo germline genetic editing technologies in arthropods, first pioneered by the Receptor-Mediated Ovary Transduction of Cargo (ReMOT Control) method. While ReMOT Control and related technologies (DIPA-CRISPR and SYNCAS) have shown great utility for gene knock-out in many different species, consistent gene insertion has been elusive, limiting widespread adoption. We developed a novel system based on the baculovirus Autographa californica multiple nucleopolyhedrovirus (AcMNPV) to deliver heritable CRISPR knock-out and knock-in constructs to the arthropod germline by oral feeding. We call this method “Germline Engineering by Viral Transduction” (“GEValT“). With GEValT, newly-eclosed female and male adult arthropods are fed on virus in sugar (and/or blood) and allowed to breed, resulting in easy generation of genetically modified offspring. GeValT is easy to use and has the potential to revolutionize gene editing methodology for researchers working across diverse arthropod species.

## Introduction

Embryonic microinjection (EM) is the primary technique used to create genetically modified arthropods. EM is a specialized technique that requires training, expertise, appropriate microinjection equipment, and is low-throughput and not scalable. In the last decade, there has been an explosion in the use of *in vivo* germline genetic editing technologies based on injection of adult arthropods, first pioneered with the development of Receptor-Mediated Ovary Transduction of Cargo (ReMOT Control). ReMOT Control relies on the evolutionarily highly conserved biological pathway of vitellogenesis to traffic molecular cargo to the developing female germline during oogenesis^1–5^. Ovary targeting peptides, derived from Yolk Protein Precursors (Ypps), are fused to cargo of interest (such as Cas9 ribonucleoprotein complex [RNP]), and when injected into the circulatory system of a female undergoing vitellogenesis, the cargo is taken up into the developing eggs^1,4^. ReMOT Control has been successfully used to genetically modify a wide range of taxa - mosquitoes and flies, beetles, wasps, kissing bugs, ticks, shrimp and prawns, and even crayfish^1–12^, and modifications of the technique such as DIPA-CRISPR^13^ and SYNCAS^14^ have shown promise in some taxa. However, while in aggregate all of these techniques are highly useful for gene knock-out, they are inefficient for knock-in of large genetic constructs.

In some systems, viral transduction vectors have proven useful as an alternative method for introducing exogenous genetic material into eukaryotic cells^15–21^. However, the utility of these viral gene delivery systems is constrained by factors such as safety (in the case where the viral vectors also infect vertebrates), virus generation, host immune responses, germline tropism, duration of transgene expression, and gene delivery packaging capacity^16–24^. Consequently, there is a significant demand for improved vector delivery technologies.

Baculoviruses constitute a family of large, arthropod-specific DNA viruses characterized by a circular dsDNA genome comprising approximately 90 to 180 genes, and predominantly act in nature as pathogens of Lepidoptera (such as the well-studied model *Autographa californica* multiple nucleopolyhedrovirus [AcMNPV]); however, they have shown plasticity in their host range^25^. In its natural lepidopteran hosts, AcMNPV infection is initiated in the midgut epithelium following ingestion of occlusion-derived virions (ODVs) released from viral occlusion bodies. After the virus traverses the midgut barrier, progeny budded virions (BVs) disseminate through the hemocoel to drive systemic infection. These two infectious forms differ in envelope composition and are specialized for oral entry (ODV) versus within-host spread (BV)^26^.

Interestingly, one study demonstrated that upon injection into the hemolymph, AcMNPV was able to infect multiple species of mosquitoes^27^. However, utility beyond somatic gene transduction was not demonstrated. Here, we show that AcMNPV can be used as an efficient transgenesis vector for arthropods, using mosquitoes as a model. Importantly, the system only works when the virus is orally fed to young individuals, and not when fed to aged individuals or when intrathoracically injected. We engineered AcMNPV to deliver Cas9 RNP and single guide RNAs (sgRNAs) (with or without cassettes for homology-directed repair insertion), demonstrate gene knock-out and knock-in, and show that the modifications are heritable. We call this technique Germline Engineering by Viral Transduction, or “GEValT”. GEValT with AcMNPV is easy to perform, eliminates the requirement for any type of injection, is scalable, relies on commercially available reagents, and is the first arthropod genetic engineering platform that allows replicable insertion of large genetic cargos without the need for EM and with no vertebrate biohazard concerns.

## Results

### AcMNPV infection in adult mosquitoes by intrathoracic injection vs. oral delivery

We investigated the temporal and spatial distribution of AcMNPV (engineered to express mCherry as a visible marker) initially using the mosquito *Ae. aegypti*, and later *Anopheles stephensi*, *An. quadrimaculatus*, and *Culex quinquefasciatus*. For these experiments, we tested both “young” (4 days post-eclosion) and “old” (15 days post-eclosion) male and female mosquitoes. For feeding treatments, we also tested both one-time infection or continuous virus exposure in the sugar. As a positive control, an identical dose of virus was injected into *Ae. aegypti* mosquitoes. Untreated mosquitoes were examined as a negative control.

After injection into *Ae. aegypti*, broad somatic infection (as measured by mCherry fluorescence) was detectable in the mosquito 3 days post-injection. Infection of the ovaries was observed, however we found that fluorescence signal in the ovaries was limited to the follicular epithelium surrounding the egg chamber, and was not present in the egg chamber itself (Figure 1).

**Figure 1.**
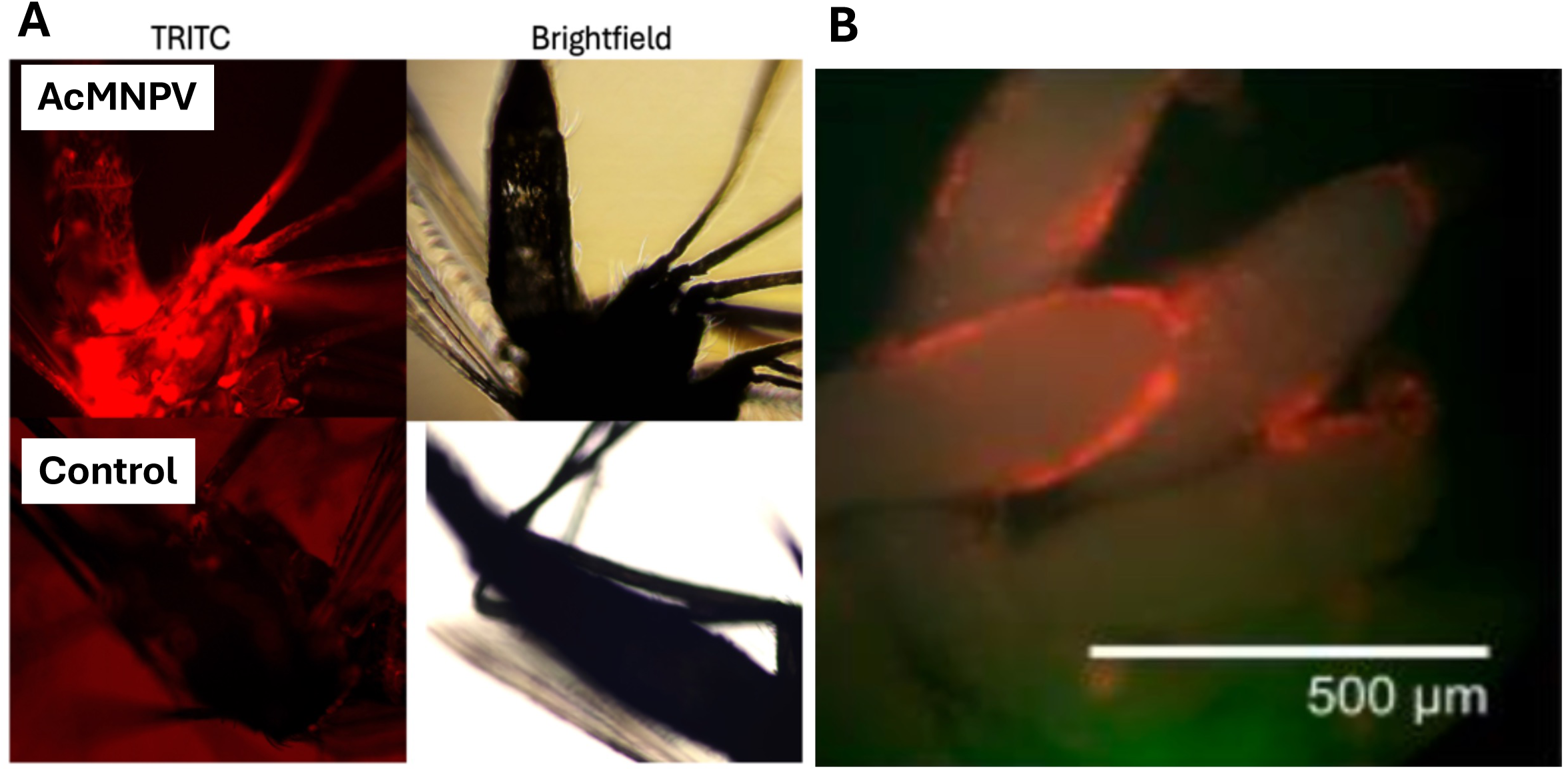
mCherry transduction in *Ae. aegypti* three days post-injection with AcMNPV. A) Whole body fluorescence (female). B) Ovary fluorescence showing follicular epithelium-localized infection.

We then tested oral delivery of AcMNPV, using *Ae. aegypti*, *An. stephensi*, *An. quadrimaculatus*, and *Culex quinquefasciatus*. After feeding on virus in the sugar, we observed high levels of viral infection/mCherry expression in the midgut and crop tissues of both male and female mosquitoes across all four species at day 3 for both tested age group (Figure 2), which disseminated to ovaries (for both age groups) and testes (in young males) within 3 days post-infection (Figure 3), but not in old males (for those species that survived past 3 days). mCherry fluorescence in ovaries and testes was ubiquitous, including in the ovarian egg chamber. After 3 days post-infection, all male *An. quadrimaculatus* and *Cx. quinquefasciatus* from the “old” treatment were deceased. Females from these species in the old treatment exhibited symptoms of illness such as lethargy, reduced feeding behavior, and an inability to fly. In contrast, “old” *An. stephensi* and *Ae. aegypti* did not exhibit obvious fitness effects, largely maintaining normal behavior and feeding activity.

**Figure 2.**
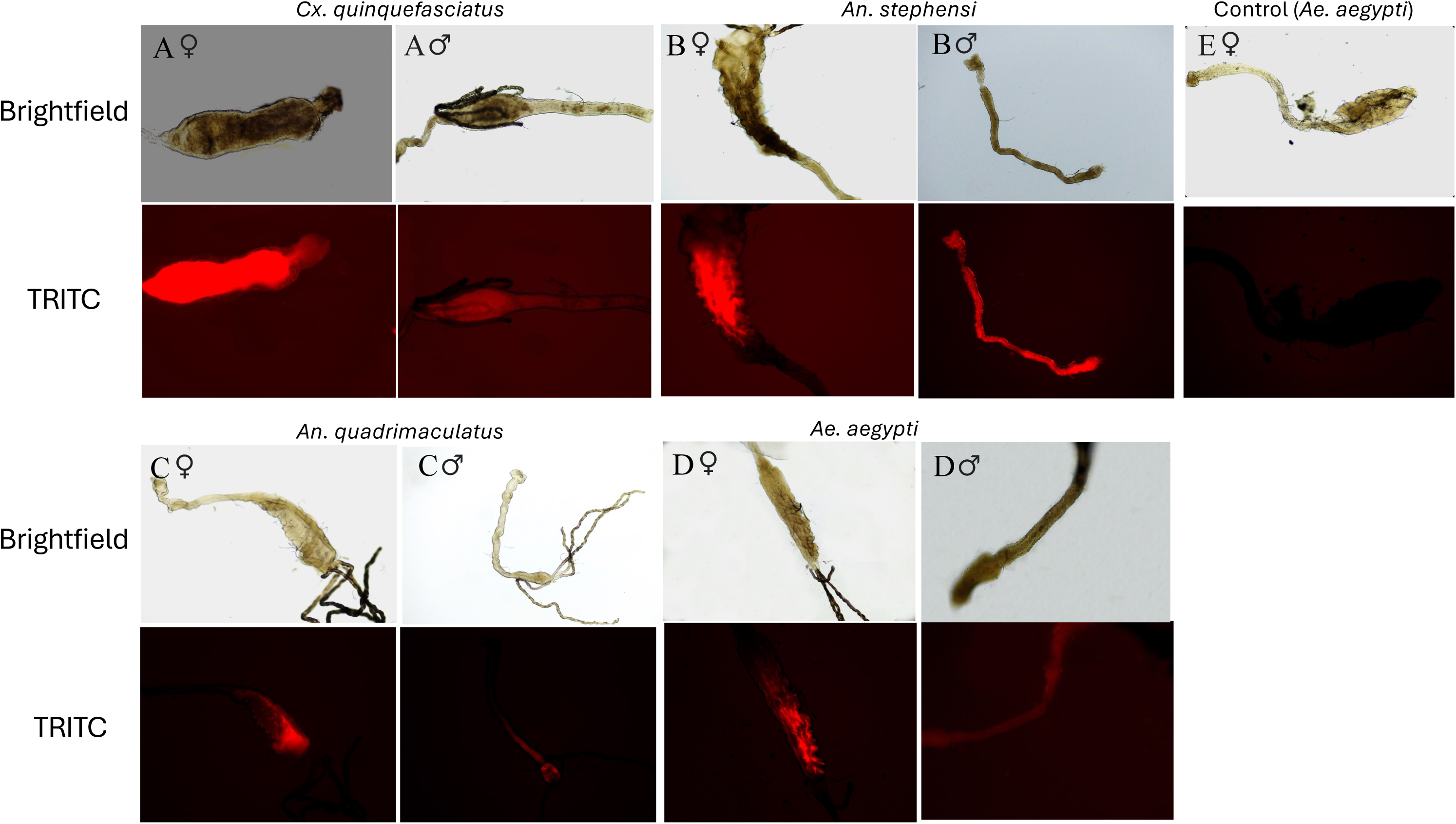
mCherry transduction in the gut of female and male A) *Cx. quinquefasciatus,* B) *An. stephensi,* C) *An. quadrimaculatus,* and D) *Ae. aegypti* female and male mosquitoes after oral exposure to AcMNPV in sugar. E is control *Ae. aegypti* female exposed to sugar without virus.

**Figure 3.**
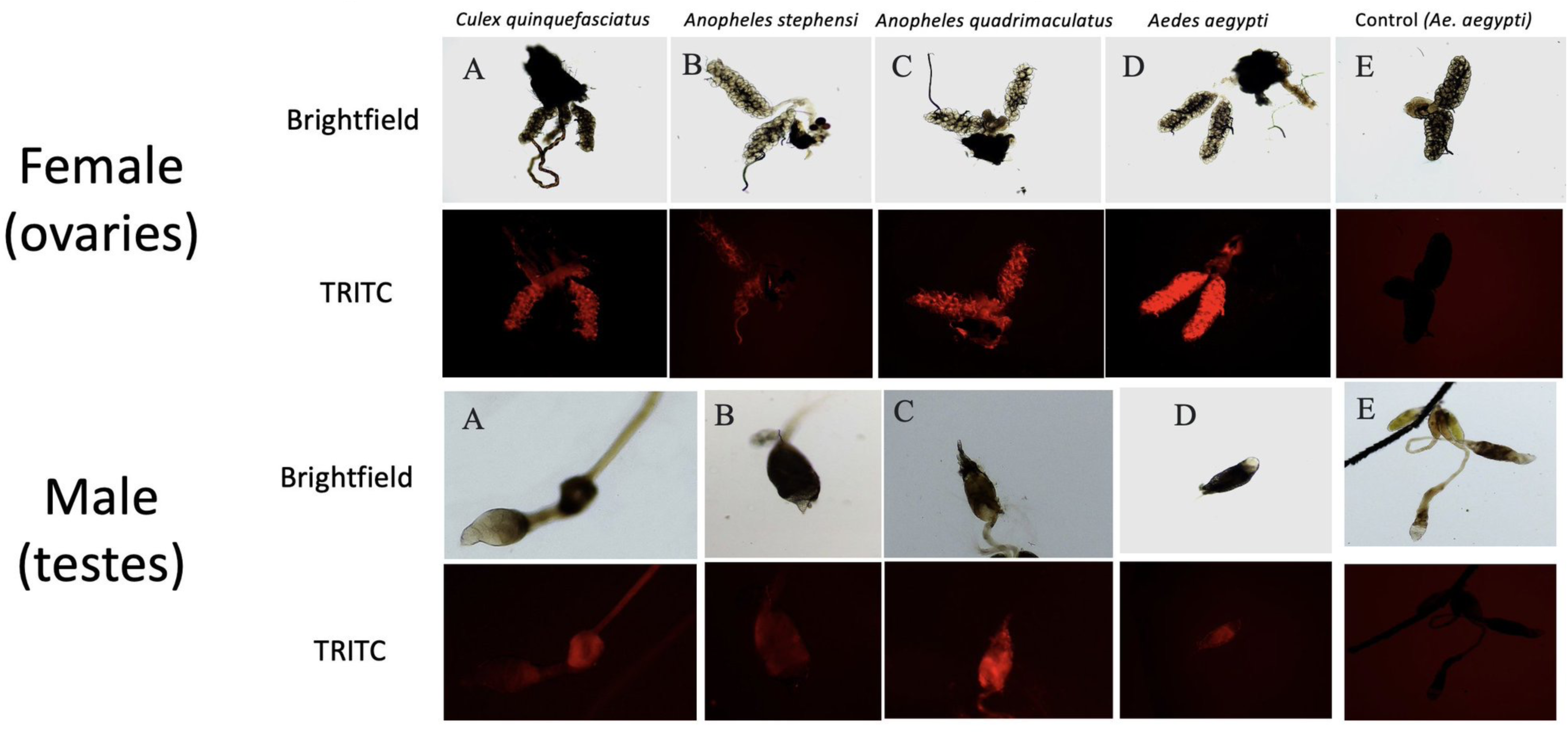
mCherry transduction in the ovaries (top) and testes (bottom) of A) *Cx. quinquefasciatus,* B) *An. stephensi,* C) *An. quadrimaculatus,* and D) *Ae. aegypti* female and male mosquitoes 3 days after oral exposure to AcMNPV in sugar. E is control *Ae. aegypti* female and male mosquitoes exposed to sugar without virus.

Using qPCR we quantified virus dynamics in gut and germline over time (*Ae. aegypti* and *An. stephensi* for a single oral viral dose, and *Ae. aegypti* for continuous oral viral exposure) in female and male mosquitoes. After a single virus dose, viral titers decreased in all tested tissues in males and females of both tested species. When continuously exposed to virus in the sugar, viral titers did not decrease in the gut of male and female *Ae. aegypti*. However, despite being exposed to virus every day, titers significantly decreased to undetectable levels in the ovaries and testes by day 10 of exposure, suggesting that virus can be maintained in the gut but is eliminated in the germline over time (Figure 4).

**Figure 4.**
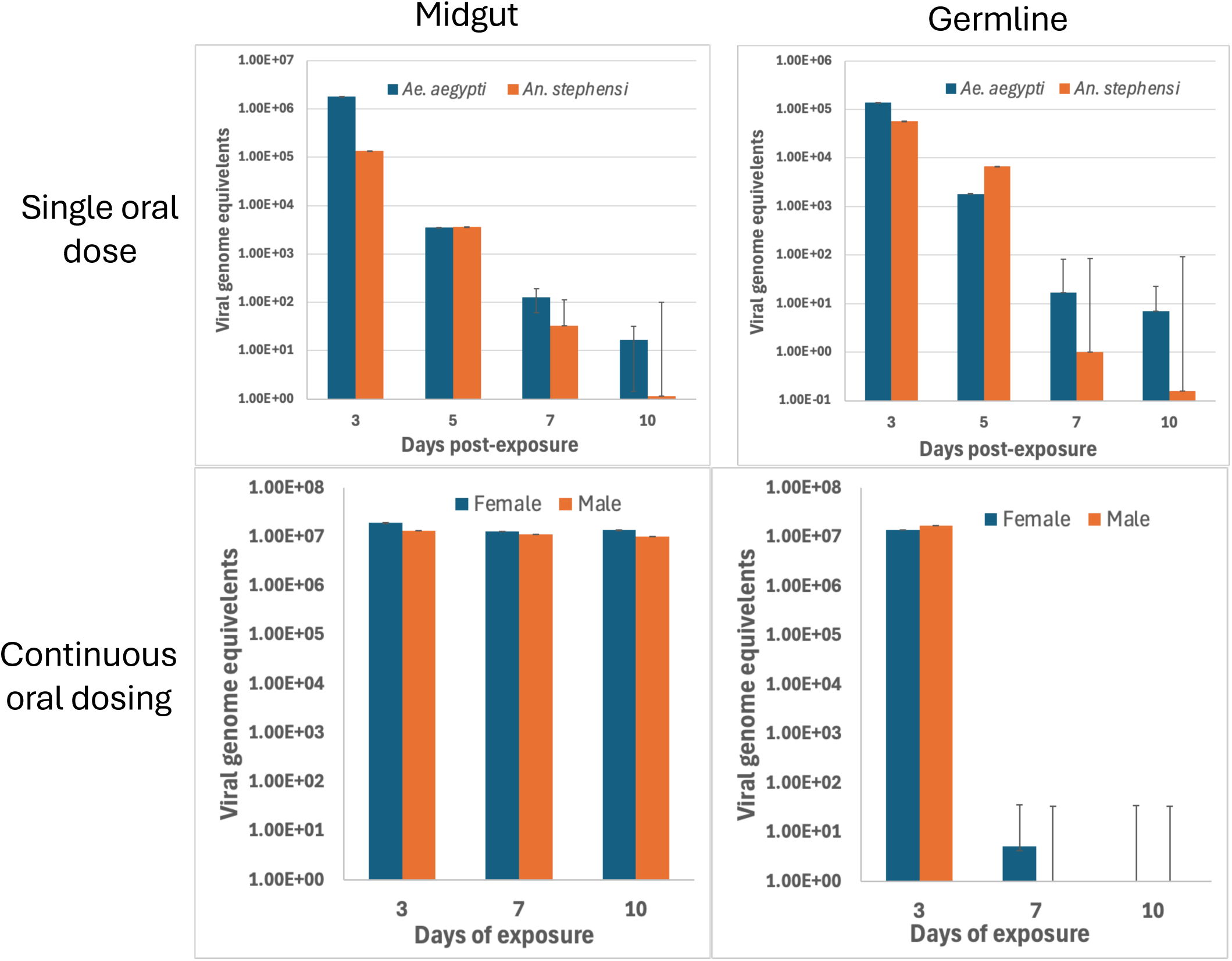
qPCR quantification of AcMNPV in mosquito tissues after oral exposure. Top: Single virus dose. AcMNPV levels over time after a single virus dose in *Ae. aegypti* and *An. stephensi*. For both species, viral levels decrease over time in both midgut and ovaries. Bottom: Continuous viral dosing. During continuous oral virus exposure, virus levels are maintained in the gut without decreasing in both male and female *Ae. aegypti*. However, virus is eliminated in the ovaries and within 7-10 days. Error bars = standard deviations.

### AcMNPV-mediated Cas9/sgRNA delivery <u>does not</u> produce knock-out phenotypes when injected into adults

Adult *Ae. aegypti* (4-5 days post-emergence) were injected with virus (10⁸–10¹⁰ FFU/mL) expressing Cas9 and either a single (460), or two (460/519) sgRNAs targeting the *kmo* gene, which causes a white eye phenotype when disrupted^1^. Four replicate experiments were conducted. No edited individuals were identified in any experiment (Table 1). These results are consistent with fluorescence data suggesting that injected AcMNPV does not reach the egg chamber in ovaries (Figure 1).

**Table 1.** *kmo* knock-out data.

| Experiment condition | Target | Viral passage | G1 offspring | Mosaic | White eye | Mutation frequency |
| --- | --- | --- | --- | --- | --- | --- |
| Control | None | N/A | NQ (>100) | 0 | 0 | 0.000 |
| Injection | kmo<br>460 | P2 | 125 | 0 | 0 | 0.000 |
| Injection | kmo<br>460 | P2 | NQ (>100) | 0 | 0 | 0.000 |
| Injection | kmo<br>460/519 | P2 | 175 | 0 | 0 | 0.000 |
| Injection | kmo<br>460/519 | P2 | NQ (>100) | 0 | 0 | 0.000 |
| Feeding 4 days old | kmo<br>460 | P2 | 196 | 0 | 0 | 0.000 |
| Feeding 4 days old | kmo<br>460 | P2 | 268 | 0 | 0 | 0.000 |
| Feeding 4 days old | kmo<br>460 | P2 | 307 | 0 | 0 | 0.000 |
| Feeding at eclosion | kmo<br>460 | P2 | 438 | **3** | **3** | **0.014** |
| Feeding at eclosion | kmo<br>460/519 | P2 | 346 | **1** | **1** | **0.006** |
| Feeding at eclosion | kmo<br>460/519 | P3 | 296 | **2** | **1** | **0.010** |

NQ = not quantified

### AcMNPV-mediated Cas9/sgRNA delivery <u>does</u> produce knockout phenotypes when fed to newly eclosed adults

Because female and male germline fluorescence levels (including visible fluorescence in the oocyte) were high in animals fed on virus, we hypothesized that oral viral exposure would be efficient at delivering CRISPR constructs to the germline. Adult *Ae. aegypti* (either newly eclosed or 4 days post-emergence) were fed virus expressing Cas9 (driven off the viral polyhedron promoter) and either a single (460), or two (460/519) sgRNAs targeting the *kmo* gene (driven off the *Ae. aegypti* U6 promoter) (Figure 4). In experimental treatments virus was added to the sugar and in the bloodmeal. Negative control treatments were fed sugar/blood without virus. No white-eye mosquitoes were identified in the control treatment, and no white-eyed phenotypes were detected among the progeny of virus-exposed 4 day old mosquitoes. However, when mosquitoes were exposed to virus in the sugar immediately upon eclosion, and given a virus-supplemented bloodmeal, white-eyed and mosaic progeny were obtained (Figure 5) (3 white-eyed pupae and 3 mosaic pupae among 438 total offspring) (Table 1). The experiment was repeated several times. In the first replicate experiment, two edited pupae (one mosaic, one fully white-eyed) were identified among 346 individuals. In the second replicate experiment, among 296 pupae, three edited individuals were detected, including one fully white-eyed and two mosaics. Mutants were obtained with virus expressing both a single sgRNA and double sgRNAs. Locus-specific PCR, cloning and sequencing of a randomly-selected white eye progeny confirmed the knock-out genotype (Figure 4). Replicate-specific editing rates between replicates ranged from 0.58% - 1.37%, with an average mutation rate of approximately 1% across all experiments using young animals (Table 1). Notably, across all experiments, gene edited white-eyed phenotypes were observed only when virus exposure began immediately after adult eclosion, and never when injected or when fed to adults 4 days post-eclosion, suggesting that the timing and route of initial virus exposure relative to adult emergence is a critical determinant of whether baculovirus-delivered editing cargo can reach the germline.

**Figure 5.**
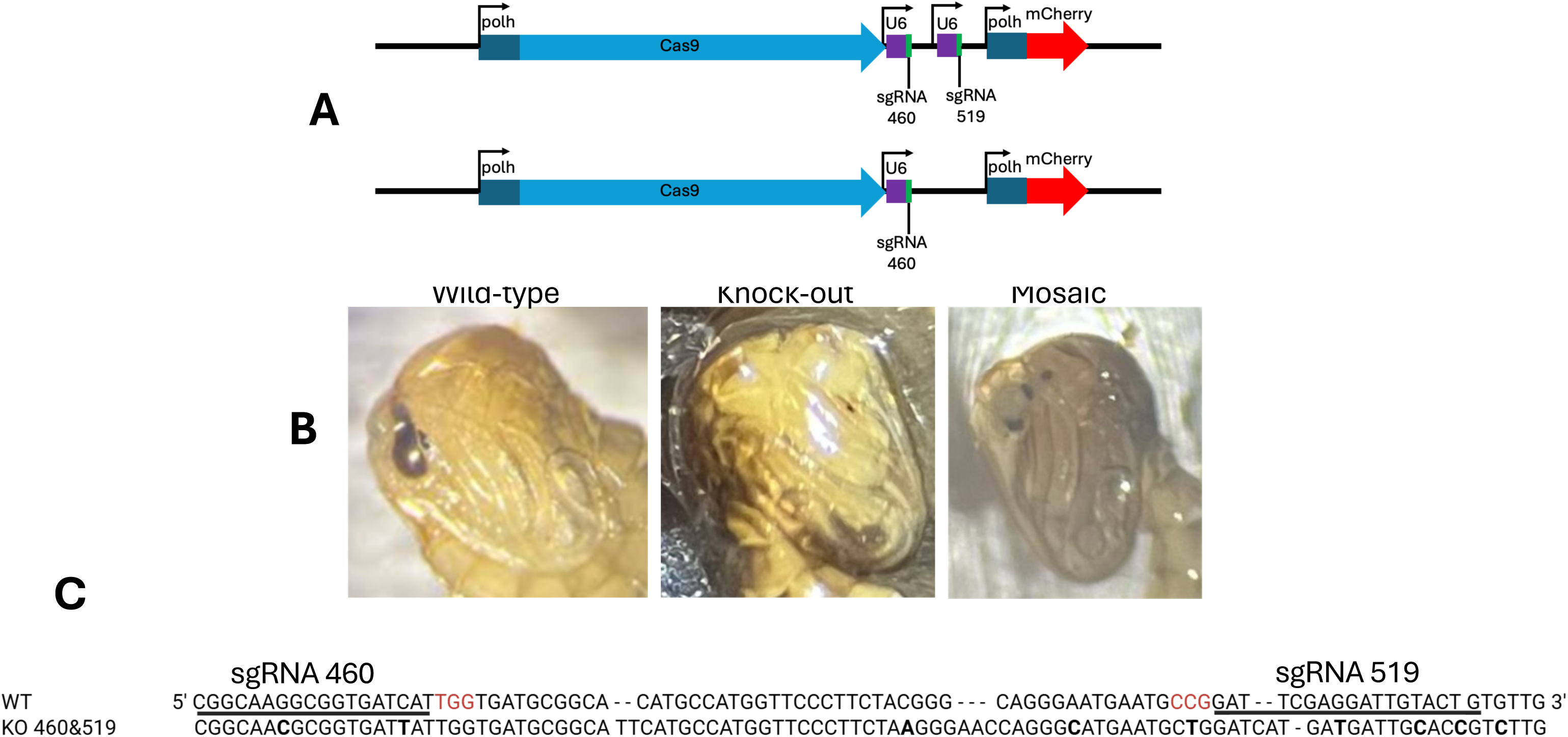
AcMNPV can generate CRISPR knock-out mutations when orally delivered. A) Schematics of AcMNPV constructs for CRISPR delivery. B) Phenotypic *kmo* mutants obtained from offspring of *Ae. aegypti* mosquitoes orally exposed to CRISPR knock-out delivery virus (wildtype, *kmo* KO, and mosaic). C) Sequence confirmation of 460/519 double mutant. sgRNA binding sites are underlined. PAM sites are delineated by red nucleotides. polh = polyhedron promoter. U6 = *Ae. aegypti* U6 promoter.

### Targeted knock-in following oral AcMNPV-mediated Cas9/sgRNA delivery

Based on the detection of sequence-confirmed knockout events, we next assessed whether a similar delivery strategy could be used for targeted knock-in by homology-directed repair (HDR). Initial plasmid constructs were designed to target the *kmo* locus by HDR; however, recombinant bacmids containing these sequences were inviable for unknown reasons. We therefore redirected the knock-in strategy to the *white* (*w*) locus, inserting mCherry as a transgenesis marker (driven off the viral polyhedron promoter). Three constructs were generated for knock-in experiments. Construct 1 comprised a pFastBac backbone carrying the left *w* homology arm, polyhedron-driven mCherry, and the right *w* homology arm. Construct 2 comprised pFastBac carrying polyhedron-driven Cas9 together with a sgRNA targeting the *w* locus. Construct 3 was an all-in-one vector combining polyhedron-Cas9, *w* sgRNA, the left *w* homology arm, polyhedron-mCherry, and the right homology arm on a single backbone (Figure 6). Newly eclosed adults were fed either Construct 3 alone, a combination of Constructs 1 and 2, or a combination of all three constructs. The regimen used in each experiment is indicated in Table 2. Because in knock-out experiments virus injection was not productive, we did not apply this technique when attempting knock-in. However, we did test whether injection could be used in concert with oral exposure to increase viral titers and potentially increase editing rates. In all cases newly-eclosed mosquitoes were used for experiments. Non-virus exposed mosquitoes were examined as a negative control.

**Figure 6.**
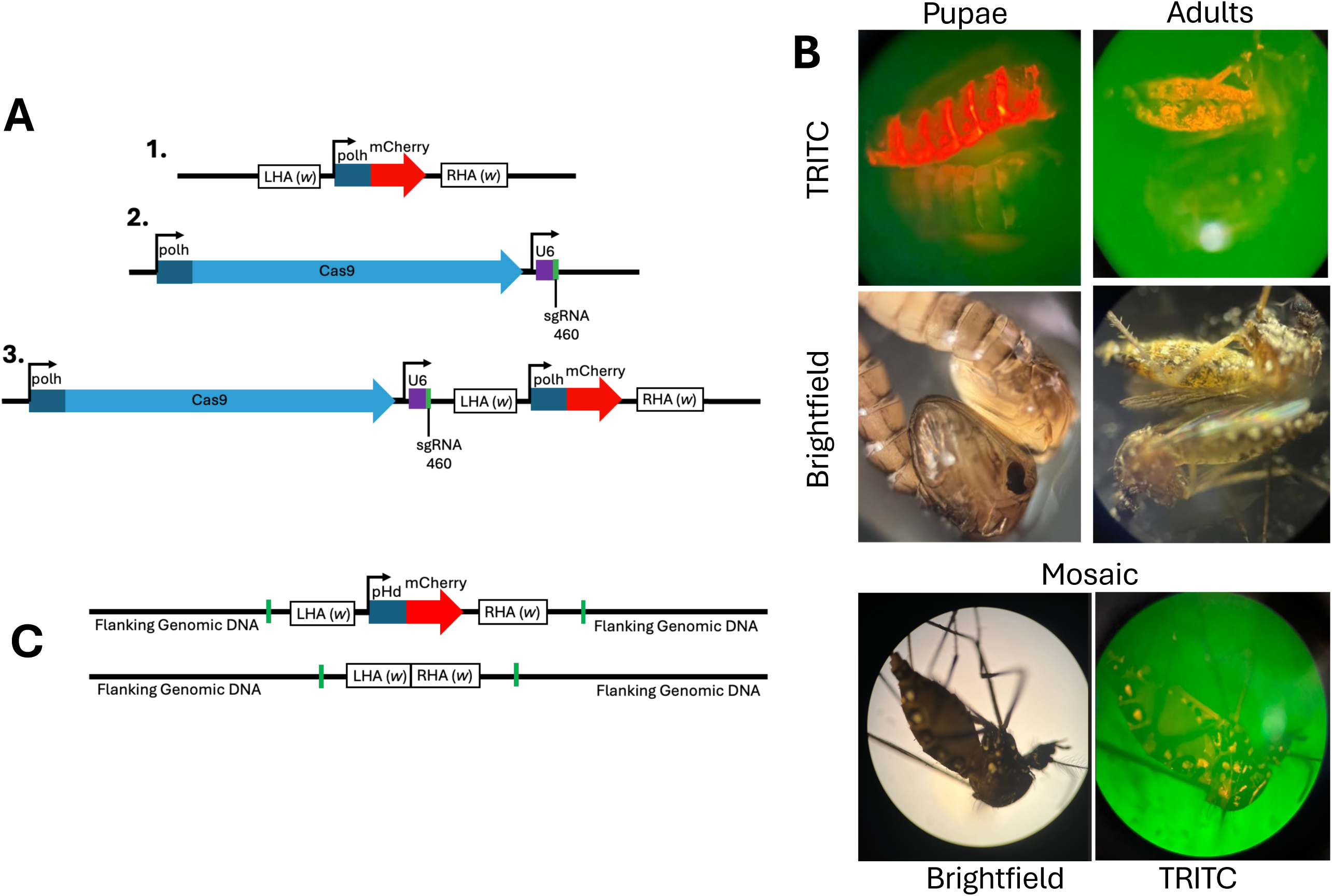
AcMNPV can generate mCherry CRISPR knock-in transgenic mosquitoes when orally delivered. A) Schematics of AcMNPV constructs 1, 2, and 3 (described in text) for CRISPR knock-in delivery by HDR into the *white* (*w*) locus. B) mCherry transgenic mosquitoes obtained from offspring of transduced *Ae. aegypti* mosquitoes (pupae, adults, and mosaic adult), and wild-type controls (pupae moved between brightfield and fluorescence images). C) Schematic of knock-in locus confirmed by sequencing. Primers in the genomic DNA flanking the insert are denoted by green lines. polh = polyhedron promoter. U6 = *Ae. aegypti* U6 promoter. LHA = Left homology arm. RHA = Right homology arm.

**Table 2.** gene knock-in data.

| Experiment condition | Construct | Viral passage | G1 offspring | Mosaic | Red | Mutation frequency |
| --- | --- | --- | --- | --- | --- | --- |
| Control | None | N/A | 296 | 0 | 0 | 0.000 |
| Feed +Injection | 1,2,3 | P2 | 0 | 0 | 0 | - |
| Feed +Injection | 1,2,3 | P3 | 8 | 0 | 0 | 0.000 |
| Feed +Injection | 1,2,3 | P4 | 0 | 0 | 0 | - |
| Feeding at eclosion | 1,2 | P2 | 358 | **1** | **2** | **0.008** |
| Feeding at eclosion | 3 | P2 | 207 | 0 | 0 | 0.000 |
| Feeding at eclosion | 1st BM: 1,2,3<br>2nd BM None | P3 | 7(1st BM),<br>115(2nd BM) | 0 | **2(1st BM)**<br>**2(2nd BM)\*** | **0.286(1st BM)**<br>**0.017(2nd BM)** |
| Feeding at eclosion | 1,2,3 | P4 | 183 | 0 | 0 | 0.000 |
| Feeding at eclosion | 1,2,3 | P4 | 45 | 0 | **4** | **0.089** |
| Breeding | None | N/A | 20 (G2) | 0 | **2** | - |

\*G1 used for heritability; BM = bloodmeal

No red individuals were observed in control mosquitoes not exposed to virus. Mosquitoes injected with virus in addition to oral exposure exhibited severe fitness deficits with very few offspring produced (Table 2), and no observable mutants. However, screening of the G1 progeny from adults exclusively orally exposed to virus (Constructs 1 and 2) identified two mCherry-positive and one mosaic among 358 pupae examined (Table 2) (Figure 6). PCR across the insert flanking regions, cloning and Nanopore sequencing confirmed locus-specific knock-in (all identified individuals were heterozygous).

In a second independent experiment, during the feeding period, 18 of 63 females died, likely reflecting toxicity at higher viral titers. The remaining 45 females in this experiment laid a total of 35 eggs, compared with 296 eggs laid by 20 non–virus-fed control females, indicating that exposure to a high virus titer significantly reduced egg production. Of 35 G1 eggs produced in the experimental group, seven hatched and developed to the pupal stage, of which two had observable mCherry expression. Two more repeats were conducted; one produced no mutants, while the other produced 4 red offspring out of 45 (Table 2). Across all successful experiments, knock-in mutants were produced when mosquitoes were exposed to a combination of constructs 1 and 2, and a combination of constructs 1, 2, and 3, but not when exposed exclusively to the all-in-one construct 3 (Table 2). Replicate-specific mutation rates ranged from 0.84% - 28.6%, with an average mutation rate of approximately 2% across all experiments using constructs 1 & 2 (with or without construct 3) (Table 2).

### AcMNPV-delivered transgene inserts are heritable

To obtain G2 progeny and confirm construct heritability, surviving G0 experimental females from above (listed in Table 2) were given a second non-infected bloodmeal and laid 150 eggs, of which 115 developed to the larval stage. Screening of these G1 larvae identified two additional mCherry-positive individuals. One mCherry-positive larva died prior to pupation; the second successfully developed to adulthood, was sib-mated, and laid 70 eggs (G2). Only 20 of these eggs successfully hatched, but of these 20 individuals two mCherry-positive animals were obtained (Table 2). Again, sequencing of the target locus confirmed targeted integration at the *w* locus. Inheritance frequencies did not confirm to expected Mendelian ratios (*X*-square 7.619, P = 0.0058), likely due to lower fitness of transgenic individuals compared to wild-type.

## Discussion

In this study, we explored the application of orally delivered AcMNPV for germline gene transduction and gene editing in arthropods, using a mosquito model. We demonstrated that when intrathoracically injected into the hemolymph of adult mosquitoes, heritable germline editing was not observed. However, when fed to newly eclosed mosquitoes (but not aged mosquitoes), AcMNPV disseminates to the germline, delivers gene editing cargo, and produces heritable targeted genome edits. We validated a *kmo* (kynurenine 3-monooxygenase) knockout by white-eye phenotypes and sequence confirmation, as well as mCherry knock-in at the *w* locus by fluorescence, sequencing of the target locus, and heritability by breeding.

A major challenge in arthropod functional genetics is that many studies still rely on embryo injections. These injections are difficult, require special skills and equipment, and are not scalable. An important advance toward bypassing embryo injections has been the development of adult-injection approaches such as ReMOT Control^1^, which can produce heritable KO edits by delivering Cas9 ribonucleoprotein complexes to adult ovaries (but not knock-in). Our findings extend the “no-embryo-injection” concept further by supporting the feasibility of an oral adult delivery route which can be used for gene knock-in, which, if generalizable, will substantially reduce barriers to adoption and enable higher-throughput experimentation^2^.

AcMNPV is a useful vector for this application because it is commercially available, can function as a high-capacity DNA delivery platform, and can efficiently transduce arthropod somatic and germline tissues. A striking observation in our experiments was that genome editing was detected only when AcMNPV exposure was oral and began shortly after eclosion, and never when the virus was injected or when mosquitoes were aged prior to exposure. This difference cannot be attributed to guide number, since editing was recovered from both single-guide and double-guide constructs, nor to the delivered titer range, which overlapped between successful and unsuccessful oral treatments. This timing dependence suggests a developmental window in the newly eclosed adult that is permissive for AcMNPV entry and subsequent germline access. One likely factor is the state of the midgut epithelium. In mosquitoes, the larval alimentary canal is nearly completely autolysed during pupation, and the adult midgut is largely built anew^28^. In the first days after emergence, the midgut epithelium is still undergoing extensive remodeling; for example Taracena-Agarwal et al.^29^ showed that approximately 20% of cells in the posterior midgut of *Ae. aegypti* incorporated nucleotide analogs within three days of eclosion, reflecting high levels of proliferation and differentiation that decline by the time the gut reaches maturity. During this period, the epithelial barrier may be more permeable to viral particles than in the mature midgut, where well-established cell junctions and a thicker basal lamina restrict paracellular and transcellular passage^30–31^. An immature barrier may allow budded virions to cross the midgut wall more readily^32^. The state of the ovaries may also be important. In newly eclosed females, the first gonotrophic cycle has not yet initiated, and ovarian follicles are in an early previtellogenic stage. If baculovirus must access oogonial stem cells or early-stage follicles to produce germline edits, a window may exist before the follicular epithelium matures and shields the developing oocyte from circulating cargo. This reasoning parallels observations from DIPA-CRISPR, a modification of the ReMOT Control method in which Cas9 ribonucleoprotein is injected into the hemocoel of adult females. Shirai et al.^33^ reported that DIPA-CRISPR was effective in *Ae. aegypti* but that the stage of the adult female at the time of injection was the single most critical parameter for editing success, and that optimal timing coincided with early oocyte development. Together, these observations suggest that both midgut permeability and ovarian accessibility contribute to the narrow temporal window we observed, and that future experiments comparing midgut barrier integrity and follicle staging between newly eclosed and 4–5 day-old adults could help identify the limiting step.

Oral infection is the natural route for baculoviruses in arthropods, with infection typically initiating in the midgut. In mosquitoes, the deltabaculovirus CuniNPV infects larval midgut epithelium following oral exposure, though its host range is restricted to *Culex* species and infection requires divalent cation supplementation; *Ae. aegypti* is not susceptible^34^. That AcMNPV transduces *Ae. aegypti* tissues following ingestion is not predicted by the natural mosquito baculovirus literature, and indicates that engineered oral delivery can access tissues not accessible to naturally mosquito-adapted baculoviruses.

Gene knock-in by HDR generally imposes stricter requirements than gene knock-out by NHEJ, including delivery of a donor sequence and engagement of a repair pathway capable of templated insertion. In mosquitoes, HDR-mediated knock-in is well documented but most often achieved via embryo-based delivery. Our finding that a baculovirus feeding route can yield sequence-verified heritable mCherry-positive knock-in alleles at the *w* locus indicates that baculovirus delivery can support templated, junction-precise integration at a defined locus. This is consistent with broader reports that baculoviral delivery can support both CRISPR knockout and donor-dependent edits in vertebrate *in vitro* systems^35^.

It is interesting that knock-in mutants were obtained after a subsequent non-infected bloodmeal in mosquitoes previously exposed to virus. Other *in vivo* transgenesis techniques such as ReMOT Control, DIPA-CRIPSR and SYNCAS preferentially edit the primary egg chamber, as this is the developing follicle that is actively undergoing vitellogenesis and thus, susceptible to uptake of the gene editing cargo^1^. In contrast, GEValT infects the entire ovary and testes, and can transduce not just the primary egg chamber but also the secondary (and perhaps tertiary etc…) egg chambers as well. Since we showed that baculovirus is eliminated from the germline within a few days post-oral exposure (Figure 4), cargo delivery and gene editing must take place rapidly across multiple stages of egg chambers simultaneously.

Our findings reinforce the need to consider marker cassette effects. A recent study in *An. gambiae* showed that when driving the marker gene DsRed off the OpIE2 promoter (derived from the baculovirus *Orgyia pseudotsugata* multicapsid nucleopolyhedrovirus [OpMNPV]), mosquitoes exhibited significant fitness effects related to bloodfeeding, reproduction, and lifespan, and that these effects were specifically related to the OpIE2 promoter and not the DsRed effector gene^36^. We saw similar significant fitness effects in generated transgenic individuals in our study (where the AcMNPV polyhedron promoter was used to drive mCherry but the mosquitoes were not infected with virus), making heritability experiments challenging due to increased mosquito mortality, markedly reduced fecundity, and feeding difficulties. To our knowledge the AcMNPV polyhedron promoter has not previously been used as a mosquito transgenesis promoter, and these results are a clear reminder that regulatory elements are not always neutral.

Several limitations of this study should be noted. Fitness effects associated with high-titer viral infection and the polyhedron promoter included reduced fecundity and survival, and bloodfeeding deficits, constraining the recovery of transgenic progeny and complicating heritability analysis. Additionally, the mechanism underlying the narrow post-eclosion window remains untested; the midgut- and ovary-based explanations proposed above are hypotheses that require direct measurement of barrier integrity and follicle staging.

Finally, there remains the question of how generalizable baculovirus-based GEValT technology will be outside mosquitoes. While we show that AcMNPV infects mosquitoes, it is canonically a lepidopteran virus. AcMNPV was first isolated from the alfalfa looper in 1969^37^, and has been (perhaps not surprisingly) shown to infect over 30 different lepidopteran species with varying degrees of pathogenicity^38^. However, a more detailed examination of published studies suggests that the AcMNPV host range is significantly more plastic than generally recognized. In addition to lepidopterans, AcMNPV can infect the insect orders Diptera (flies)^27,38–40^, Coleoptera (beetles)^41^, Hymenoptera (bees/wasps/ants)^42^, Blattodea (cockroaches)^40^, and even non-insect arthropods such as ticks^43^. These data suggest that genetic modification by GEValT may be broadly generalizable across diverse arthropod taxa. It should also be noted that GEValT, as a conceptual idea, is not limited to transduction with AcMNPV. Insect-specific viruses (ISVs) are ubiquitous across arthropods where they are maintained by infection of the germline and subsequent vertical transmission^20–21,44^. These ISVs could in principle be isolated and engineered as GEValT vectors for species recalcitrant to baculovirus infection. Additionally, if viruses with the correct germline tropism properties were identified, GEValT could in concept even be extended to vertebrate taxa.

## Material and Methods

### Cells and viruses

The Bac-to-Bac baculovirus expression system (Thermo Fisher Scientific) was employed for generation of recombinant baculovirus. A codon-optimized mCherry coding sequence was synthesized as a gBlock gene fragment (Integrated DNA Technologies) and cloned into pJET1.2/blunt (Thermo Fisher Scientific, cat. 10359-016)) to generate pJET-mCherry. The mCherry cassette was then PCR-amplified from pJET-mCherry and subcloned downstream of the polyhedron (*polh*) promoter to generate the final transfer vector, which was recombined into the bacmid and used for transfection of Sf9 cells. Viral stocks were stored at −80 °C in the respective culture media.

### Generation of viral passage stocks

Recombinant bacmid DNA was transfected into Sf9 cells using the Bac-to-Bac Baculovirus Expression System (Thermo Fisher Scientific, cat. 10359-016) according to the manufacturer’s instructions. Transfected cells were inspected daily from 72 h post-transfection for signs of late-stage infection. Medium was harvested at 72–96 h post-transfection, clarified by centrifugation at 500 × g for 5 min, and retained as the P0 stock. Because P0 stocks are low in both titer (typically 10⁶–10⁷ FFU/mL) and volume, P0 was used to infect Sf9 cells seeded at 2 × 10⁶ cells/mL at an MOI of 0.05–0.1, which were incubated at 27 °C for 72–96 h to generate the P1 stock. The same procedure was repeated at increasing scale to generate P2, P3, and P4 stocks. Titers were determined by focus-forming assay (FFA) and ranged from 10⁸–10¹⁰ FFU/mL. Working stocks were stored at 4 °C protected from light, with aliquots archived at −80 °C.

### Knock-out constructs

Two sgRNAs targeting the genetic locus at nucleotides 460 (sgRNA460) and 519 (sgRNA519) of exon 5 of *kmo* previously used by Chaverra-Rodriguez et al.^1^ was used for the knockout and knock in experiment. The two primers within each pair exhibit complementarity over the 20 nucleotides specific to the chosen gRNA target, and their annealing results in a linker with specific overhangs designed for the two *BbsI* sites present in the cloning vectors. U6::gRNA constructs were created by individually cloning gRNA target DNA sequences 1, and 2 into the *BbsI* site of two separate pKSB-sgRNA (Addgene :pKSB-609 U6-sgRNA1 ; pKSB-610 U6-sgRNA2 cat no. 173671 and 173672) vectors followed by confirmation through sequencing. The confirmed two pKSB-*kmo1*-gRNA1, pKSB-*kmo1*-gRNA2, along with mCherry and Cas9 (add gene: pDSAY-vasaCas9sv40 cat no. 173669) were assembled into the pFastBac vector from Bac-to-Bac baculovirus expression system (Thermo Fisher Scientific cat no. 10359016) using NEBuilder® HiFi DNA Assembly Master Mix (cat no. E2621L). The confirmed pFastBac vector transformed into MAX Efficiency™ DH10Bac Competent Cells. The confirmed rbacmid then was used to infect sf9 cells.

### Knock-in constructs

Cas9 ribonucleoprotein components were designed with sgRNAs targeting *kmo* and the *white* gene (*w*; AAEL016999), with the *w* sgRNA targeting exon 3 (Aeag white sgRNA1^45^; (Supplementary Table 1). HDR donor templates were designed to mediate site-specific integration of mCherry at the *w* locus, using locus-matched 5′ and 3′ homology arms. For the *w* knock-in workflow, three transfer constructs were assembled: (i) a Cas9/sgRNA expression plasmid (polh– Cas9 with an sgRNA targeting *w*), (ii) a donor plasmid carrying the repair template (5′ homology arm–polh-mCherry–3′ homology arm), and (iii) an all-in-one plasmid combining the Cas9/sgRNA cassette and HDR donor template on a single vector (–polh–Cas9/sgRNA[*w*]–5′ homology arm– polh-mCherry–3′ homology arm). Constructs were assembled using NEB HiFi DNA Assembly Master Mix with fragments designed in NEBBuilder. The assembled CAS9/gRNA and homology arms were subsequently integrated into the pFastBac vector from the bac-to-bac expression system (Thermo fisher; Cat NO10359016). The verified vector was then transformed into MAX Efficiency™ DH10Bac Competent Cells. After verification of the recombinant bacmid sequence, it was used to infect Sf9 cells.

### Cell culture

*Spodoptera frugiperda* SF9 cells (Thermo Fisher Scientific) were grown at in monolayers at 27°C in Sf-900™ III SFM insect medium (Catalogue number 12658019; ThermoFisher) supplemented with sodium bicarbonate (0.35 g/L), 10% fetal bovine serum (FBS) (35-010-CV, Coring), and a 1% solution of Penicillin-Streptomycin (5,000 U/mL) (Catalogue number 15070063, Gibco).

### Mosquito rearing

*Ae. aegypti*, *An. stephensi*, *An. quadrimaculatus*, and *Cx. quinquefasciatus* were reared at 27 °C and at 80% humidity under a 12-h light/dark cycle. Larval stages were fed on pulverized fish food. Adults were provided with 5% glucose through a cotton wick.

### Focus Forming Assay (FFA)

Recombinant baculovirus titer was determined by focus-forming assay based on mCherry expression. One day prior to infection, 96-well plates were seeded with 1-2 × 10^5^ Sf9 cells per well in 100 μL of growth medium to establish a confluent monolayer. Ten-fold serial dilutions of virus were prepared in FBS-free medium. Growth medium was removed from the seeded plates, and cells were infected with 30 μL of each virus dilution per well and incubated at 27°C for 1 hour to allow viral entry. The inoculum was then aspirated and replaced with 70-100 μL of complete growth medium containing 0.8% methylcellulose, and plates at 27°C for 48 hours to focus formation. The overlay medium was removed, and cells are fixed with 50 μL of 4% formaldehyde in PBS for 30 minutes at room temperature and washed with PBS. Foci were visualized and counted by mCherry fluorescence on a fluorescence microscope using a filter set appropriate for mCherry (ex ∼587 nm/em ∼610 nm), and titers were calculated as focus-forming units per mL (FFU/mL).

### Baculovirus transduction and tissue tropism experiments

To characterize the spatial and temporal distribution of baculovirus in adult mosquitoes, young (4 days post-eclosion) and old (14 days post-eclosion) male and female *Ae. aegypti*, *An. stephensi*, *An. quadrimaculatus*, and *Cx. quinquefasciatus* were exposed to baculovirus through sugar feeding. Viral inoculum was prepared by dissolving 0.10 g sucrose (Sigma Life Sciences, S0389) in 1.0 mL virus stock (final concentration 10% w/v). Cotton balls saturated with virus-sucrose solution were placed on top of cages and replaced daily. Mosquitoes were injected or exposed to continuous virus feeding at a dose of passage 2 (P2; 10⁷–10¹⁰ FFU/mL). Untreated mosquitoes served as a negative control. Mosquitoes were assessed for baculovirus infection by fluorescence microscopy and qPCR with midgut, ovary, and testes examined separately.

### Baculovirus-mediated genome editing: oral delivery

For knock-out and knock-in editing experiments, two exposure timing conditions were tested. In the first condition, adult *Ae. aegypti* (Liverpool) were maintained on control sucrose (no virus) for 4–5 days post-eclosion before receiving a virus-containing blood meal. In the second condition, newly eclosed adult mosquitoes were provided virus-sucrose solution beginning within 12 hours of adult emergence and fed continuously for 4–5 days, after which a virus-containing blood meal was administered. In both conditions, viral inoculum was prepared daily as described above.

Depending on the experiment, passage 2 (P2; 10⁷–10¹⁰ FFU/mL), passage 3 (P3; 10¹⁰ FFU/mL), or passage 4 (P4; 10⁹ FFU/mL) virus was used. To ensure Cas9 expression from the constructs, qPCR was conducted measuring Cas9 transcript levels in P1 and P2 stocks. A minimum of 55 females and 35 males was used per cage per experiment. Three days after blood feeding, an egg cup was placed in the cage and eggs were collected. Resulting offspring were screened for white-eye phenotypes (knockout) or mCherry fluorescence (knock-in). Control cages received no virus in either sugar or blood meals.

### Baculovirus-mediated genome editing: injection delivery

Virus was delivered by direct intrathoracic microinjection of 0.69 nL of P2 virus (10⁸–10¹⁰ FFU/mL). For knock-out experiments, mosquitoes were only exposed to virus by injection. For knock-in experiments, injection was instead applied as a supplement to the oral regimen to test whether direct introduction of virus into the hemocoel would increase germline delivery beyond that achieved by feeding alone. In this case, newly eclosed adults were provided virus-supplemented sucrose and fed continuously for 4–5 days, followed by a virus-supplemented blood meal. One day after bloodfeeding, females were injected intrathoracically with 0.69 nL of virus (regimen indicated in Table 2) Offspring in both experiments were screened as described above.

### Molecular confirmation of genome editing events

Genomic DNA was extracted from individual mosaic and white-eyed pupae (knock-out) or mCherry-positive individuals (knock-in) using the E.Z.N.A. Tissue DNA Kit (Omega) according to the manufacturer’s suggested protocol. For knockout confirmation, the *kmo* target region spanning sgRNA sites 460 and 519 was PCR-amplified using primers flanking the expected cut sites (Supplementary Table 1). For knock-in confirmation at the *w* locus, primers annealing outside the 5′ and 3′ homology arms were used to amplify across the full integration site (the “Arm to Arm primer outside the construct” listed in Supplementary Table 1). PCR products were separated by 1% agarose gel electrophoresis, and bands of the expected size were excised and gel purified. Purified fragments were cloned into the pJET1.2/blunt cloning vector (Thermo Fisher Scientific) following the manufacturer’s protocol. Plasmid DNA from individual colonies was submitted for Oxford Nanopore long-read sequencing (Plasmidsaurus). Resulting reads were aligned to the reference sequences to confirm indels at the *kmo* locus (knockout) or the expected junction structure at the *w* locus (knock-in).

### Quantification of baculovirus DNA by real-time qPCR

Ovary, testes and midgut for both female and male mosquitoes were washed three-times with 1X PBS remove free virus particles from the organ. Total cellular DNA was extracted using the E.Z.N.A tissue DNA kit (Omega). The quantity and quality of DNA was checked using a NanoDrop ND-1000 spectrophotometer (Thermo Fisher Scientific). Real time-qPCR was performed in a 386-well plate, with each well containing 2 μl DNA (100 ng/µl), 1 μl 10 μM specific primers (final concentration, 300 nM), 6 μl H2O, and 10 μl Universal SYBR Green super Mix (Biorad). Samples were run in three biological triplicates against a standard curve generated from bacmid plasmid DNA. Primers are listed in Supplementary Table 1.

## AI statement

No generative AI was used in the writing of this manuscript.

## Acknowledgements

We thank Francine McCullough and Sage McKeand for assistance with laboratory experiments.

## Funding

This research was supported by NIH grant R01AI128201, NSF grant 2525447, USDA Hatch Project 7011141, bridge funds from the Penn State Huck Institutes of the Life Sciences and the Penn State College of Agricultural Sciences, and funds from the Dorothy Foehr Huck and J. Lloyd Huck endowment to JLR.

## Competing interests

The authors have filed for provisional patent protection on the GEValT technology.

---

**Supplementary Table 1.**
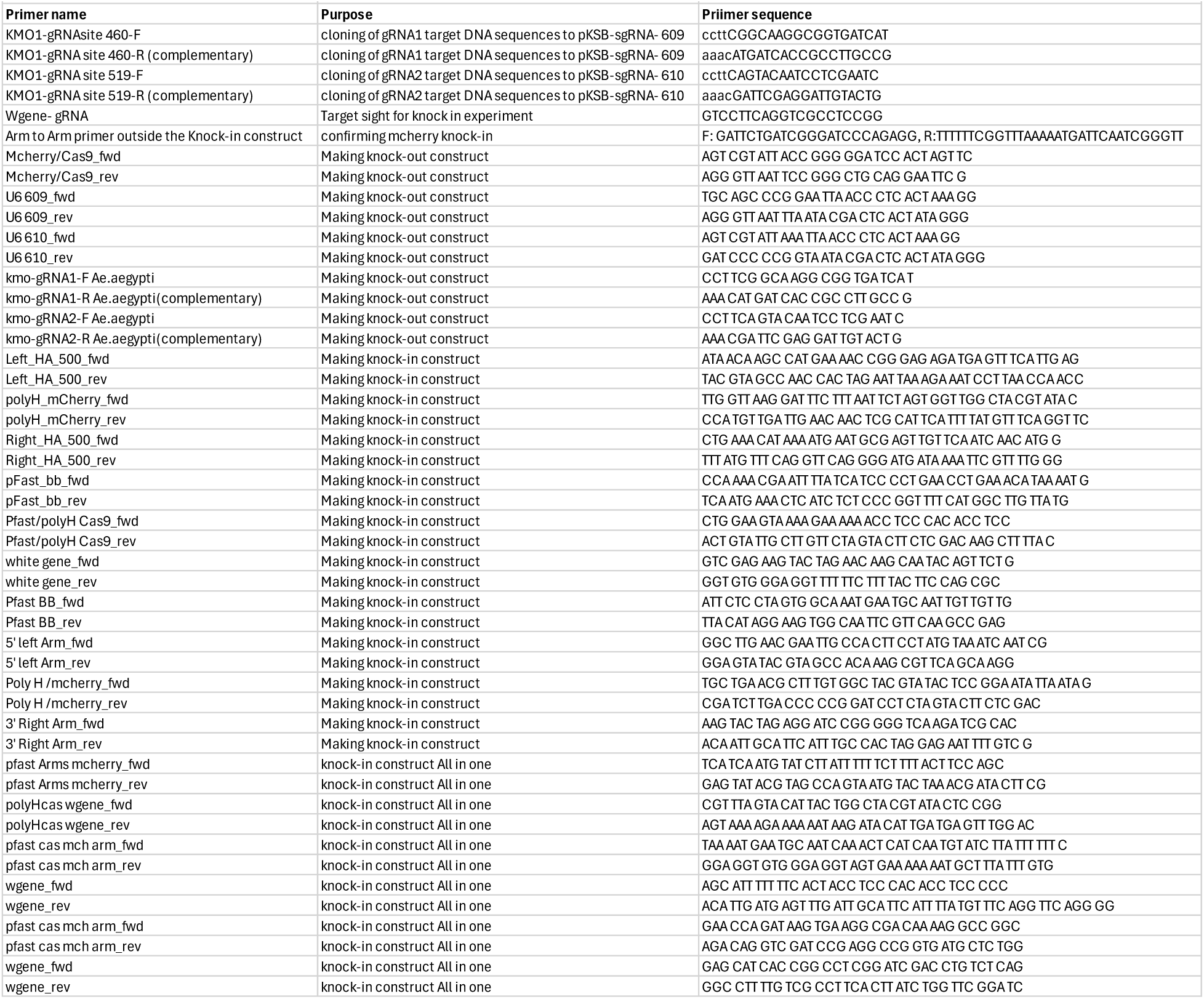
Oligonucleotide sequences used in this study.

